# CTCF-binding sites demarcate chromatin domains enriched with H3K9me2 or H3K9me3 and restrict the spreading of these histone modifications in human cells

**DOI:** 10.1101/2025.05.28.656735

**Authors:** Kyung-Ju Shin, Jin Kang, AeRi Kim

## Abstract

CTCF-binding sites are frequently found at the boundaries of chromatin domains enriched with specific histone modifications in vertebrate genomes, where they function as barriers that limit the spreading of these modifications. However, their role in domains marked by H3K9me2 or H3K9me3 remains poorly understood. In this study, we analyzed CTCF-binding sites and the distribution of H3K9me2 and H3K9me3 around them in human K562 cells, and investigated the spreading of these modifications following CTCF depletion. Genome-wide analysis revealed that certain CTCF-binding sites are located at the boundaries of chromatin domains enriched with H3K9me2 or H3K9me3. Upon loss of CTCF binding, both modifications spread to neighboring chromatin at over half of the boundaries delimiting highly and distinctly enriched domains. Boundaries where such spreading did not occur were flanked by chromatin in a more active state compared to those where spreading occurred. Gene transcription was significantly reduced in neighboring regions where H3K9me2 or H3K9me3 had spread. Further comprehensive analysis revealed that this transcriptional reduction was specifically associated with the spread of H3K9me2. These findings suggest that CTCF-binding sites demarcate chromatin domains enriched with H3K9me2 or H3K9me3 and act as barriers to restrict the spread of these modifications in mammalian cells. Notably, the spread of H3K9me2 appears to play a specific role in transcriptional repression.

## Introduction

CTCF is a DNA-binding protein whose binding sites function as insulators that block the transmission of enhancer activity to non-target promoters and/or demarcate chromatin domains in vertebrate genomes (Hou et al. 2008; Kim et al. 2015; Ali et al. 2016). Genome-wide mapping across various human cell types has shown that CTCF-binding sites are enriched at the boundaries of chromatin domains marked by histone H3K27me2 (Cuddapah et al. 2009). Loss of CTCF binding at insulator sequences is associated with the expansion of H3K27me3 into tumor suppressor genes such as the *RASSF1* in cancer cells (Chang et al. 2010). In the *HOXA* locus, specific CTCF-binding sites are located between two distinct chromatin domains marked by H3K4me3 and H3K27me3, respectively, and upon the loss of CTCF-binding one domain expands into the other, accompanied by the spreading of histone modifications (Kim et al. 2011a; Narendra et al. 2015). Our previous study showed that a H3K27ac- enriched domain, including the transcriptionally active *beta-globin* locus, expanded into adjacent H3K27me3-enriched neighboring region following the deletion of multiple CTCF sites encompassing the domain (Kang et al. 2021). Collectively, these findings suggest that CTCF-binding sites delimit chromatin domains enriched with specific histone modifications, block the spreading of histone modifications and, in turn, protect domains from encroachment by neighboring domains. These roles of CTCF-binding sites may contribute to the establishment and maintenance of chromatin environments conductive to proper gene transcription.

Two forms of methylation at histone H3K9, H3K9me2 and H3K9me3, are enriched in heterochromatin, which exhibits a more compact and transcriptionally repressive structure compared to euchromatin. The initiation and spreading of these modifications during heterochromatin assembly have been extensively studied in fission yeast. These processes are mediated by the recruitment of histone methyltransferases and are reinforced through positive feedback loops involving these enzymes and chromatin binding proteins, such as HP1, which recognize H3K9 methylation (Allshire and Madhani 2018; Grewal 2023). An early study in fission yeast showed that two inverted repeats within the *MATA* locus act as boundary elements that prevent the spreading of H3K9me3-enriched heterochromatin (Noma et al. 2001). Spreading of H3K9me2 and H3K9me3 has been observed in mammalian cells. For instance, silencing of the *DNTT* gene during stimulated thymocyte maturation is accompanied by H3K9me2/3 enrichment at its promoter, which then spreads both upstream and downstream (Su et al. 2004). Similarly, during lineage commitment and differentiation of hematopoietic stem cells, H3K9me2 peaks arise in genic regions and expand to form large H3K9me2 domains (Chen et al. 2012). Notably, the spreading of H3K9me2 and/or H3K9me3 has been suggested to be associated with the loss of CTCF binding in the *DMPK* gene with expanded CTG repeats in myotonic dystrophy (Cho et al. 2005), and in human bronchial epithelial cells exposed to nickel (Jose et al. 2014).

H3K9me2 and H3K9me3 are closely associated with gene repression. A large-scale genome-wide study revealed that these modifications are more enriched in transcriptionally inactive genes compared to active ones, and their levels near transcription start sites (TSSs) of genes are negatively correlated with gene transcription levels (Barski et al. 2007). In mouse embryonic fibroblasts, genes with H3K9me2- or H3K9me3-marked promoters are more strongly repressed than those lacking these modifications, and this repression is alleviated upon depletion of the histone methyltransferases responsible for their deposition (Montavon et al. 2021). In embryonic stem cells and preadipocytes, the transcriptional repression of developmental genes and the adipogenesis master regulator gene *PPARG*, respectively, is mediated by H3K9me2 (Tachibana et al. 2002; Wang et al. 2013). Transcriptional silencing during cell maturation and differentiation is frequently accompanied by the acquisition of H3K9me2 or H3K9me3 (Su et al. 2004; Chen et al. 2012). These findings collectively suggest that both H3K9me2 and H3K9me3 contribute to the repression of gene transcription. However, it is unclear whether these two modifications are deposited together in genomic regions. In mammalian cells, H3K9me2 and H3K9me3 are primarily catalyzed by different histone methyltransferases (Rice et al. 2003; Montavon et al. 2021), and they have been reported to localize to distinct repressive chromatin by immunofluorescence staining (Rice et al. 2003).

CTCF binding appears to play a critical and direct role in demarcating chromatin domains enriched with specific histone modifications, such as H3K27me2, H3K27me3, H3K4me3, and H3K27ac, and in preventing the spreading of these modifications, as demonstrated at several genomic loci (Cuddapah et al. 2009; Chang et al. 2010; Kim et al. 2011a; Narendra et al. 2015; Kang et al. 2021). However, the role of CTCF binding in chromatin domains enriched with H3K9me2 or H3K9me3 remains unclear. In this study, we conducted a genome-wide analysis of CTCF-binding sites in human K562 cells and examined the distribution of H3K9me2 and H3K9me3 around these sites. We identified chromatin domains that were demarcated by CTCF-binding sites and were highly and distinctly enriched with either H3K9me2 or H3K9me3. To investigate the role of CTCF binding in these domains, we depleted CTCF protein and assessed changes in histone modifications in neighboring chromatin regions. In addition, gene transcription was analyzed in the neighboring regions upon the spreading of histone modifications. Obtained results suggest that a subset of CTCF-binding sites function as boundaries of chromatin domains enriched with H3K9me2 or H3K9me3 and also act as barriers that restrict the spreading of these modifications. This spreading appears to be influenced by the chromatin landscape of neighboring regions. Notably, the spreading of H3K9me2 is specifically associated with transcriptional repression of genes.

## Results

### CTCF-binding sites demarcate chromatin domains enriched with H3K9me2 or H3K9me3

To investigate whether CTCF-binding sites are located at the boundaries of domains enriched with H3K9me2 or H3K9me3 at the genome-wide level, we performed ChIP-seq analysis in control K562 cells, which served as the baseline for the CTCF depletion experiments in this study. ChIP-seq data from two biological replicates were compared each other in each experimental condition and then merged prior to downstream analysis (Supplemental Fig. S1). A total of 46,927 CTCF-binding sites were identified by genome-wide mapping (Fig. 1A). ChIP-seq signals for H3K9me2 and H3K9me3 were analyzed within 20 kb regions upstream and downstream of these CTCF-binding sites (Fig. 1B). Signal alignment based on the differential enrichment between the upstream and downstream regions revealed sharp transition in histone modification levels at a subset of CTCF-binding sites. For H3K9me2, signal intensity was significantly higher in the half of the regions (upstream or downstream) where the difference exceeded 120, compared to the other half (Fig. 1C). A similar pattern was observed for H3K9me3, with signal differences exceeding 100 (Fig. 1D). Notably, CTCF binding was stronger at sites exhibiting greater histone modification differences. These findings suggest that a subset of CTCF-binding sites may function as boundaries demarcating chromatin domains enriched in H3K9me2 or H3K9me3, and that this boundary function is associated with robust CTCF binding.

**Figure 1.**
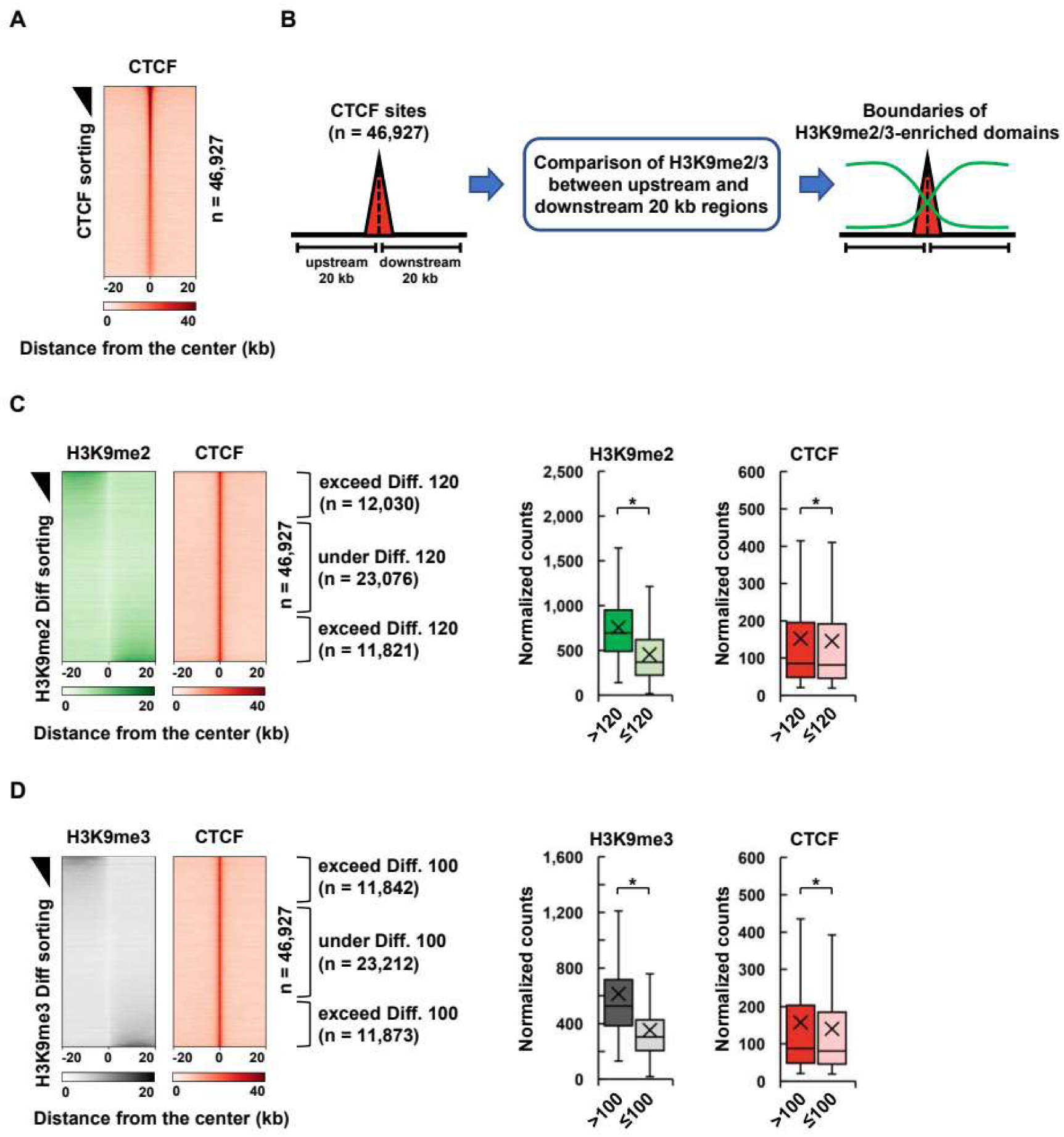
Histone H3K9me2 and H3K9me3 around CTCF-binding sites in human K562 cells. (A) ChIP-seq signals of CTCF around CTCF-binding sites (n = 46,927) were visualized as a heatmap in control K562 cells, which served as the baseline for the CTCF depletion experiments in further study. The sites were sorted based on CTCF binding intensity. (B) Schematic diagram illustrates the comparison of H3K9me2 or H3K9me3 signals between 20 kb upstream and downstream regions of CTCF-binding sites. (C) H3K9me2 and CTCF signals around CTCF-binding sites were depicted in heatmaps. The sites were sorted based on the H3K9me2 signal differences (Diff). Normalized counts of H3K9me2 and CTCF were measured within the 20 kb regions where the H3K9me2 Diff exceeded 120 (n = 23,851) or did not (n = 23,076), and at the corresponding CTCF-binding sites, respectively. (D) CTCF-binding sites were sorted based on H3K9me3 Diff. Normalized counts of H3K9me3 and CTCF were measured within 20 kb regions showing a Diff greater than 100 (n = 23,715) or not (n = 23,212), and at the corresponding CTCF sites, respectively. P-values were calculated using the Mann-Whitney U test. *P < 0.01

### H3K9me2 and H3K9me3 spread from their enriched domains into neighboring chromatin upon CTCF depletion

To explore the spreading of H3K9me2 and H3K9me3 upon the loss of CTCF binding, we inhibited the expression of CTCF gene in K562 cells using shRNA. A pronounced reduction was detected at both mRNA and protein levels by day 3 after lentiviral transduction, and this reduction was sustained through day 9 (Fig. 2A). Consequently, ChIP-seq analysis revealed a substantial loss of CTCF binding at its target sites by day 6 in CTCF knockdown (CTCFi) cells compared to control K562 cells (Fig. 2B). To identify chromatin domains that are demarcated by CTCF-binding sites and are highly and distinctively enriched with H3K9me2, we compared ChIP-seq signals within 20 kb regions upstream and downstream of these sites. A total of 7,195 H3K9me2-enriched regions were defined based on a signal difference exceeding 100 and a greater than 2-fold change (Fig. 2C). In CTCFi cells, H3K9me2 signals significantly increased in neighboring regions flanking these enriched domains. Specifically, 4,436 neighboring regions (61.7%) showed increased H3K9me2 levels, while 2,759 regions (38.3%) showed no increase (Fig. 2D). Using the same criteria-signal difference greater than 200 and a more than 2-fold change, we identified 7,887 H3K9me3-enriched regions (Fig. 2E). Similar to H3K9me2, H3K9me3 signals significantly increased in adjacent regions in CTCFi cells, with 5,262 neighboring regions (66.7%) exhibiting elevated H3K9me3 levels and 2,625 regions (33.3%) showing no elevation (Fig. 2F). CTCF binding was similarly reduced, irrespective of the increase in these modifications in neighboring regions. Collectively, these results show that CTCF depletion facilitates the spreading of H3K9me2 and H3K9me3 from their enriched domains into neighboring chromatin, supporting a key role for CTCF in restricting the propagation of these repressive histone modifications.

**Figure 2.**
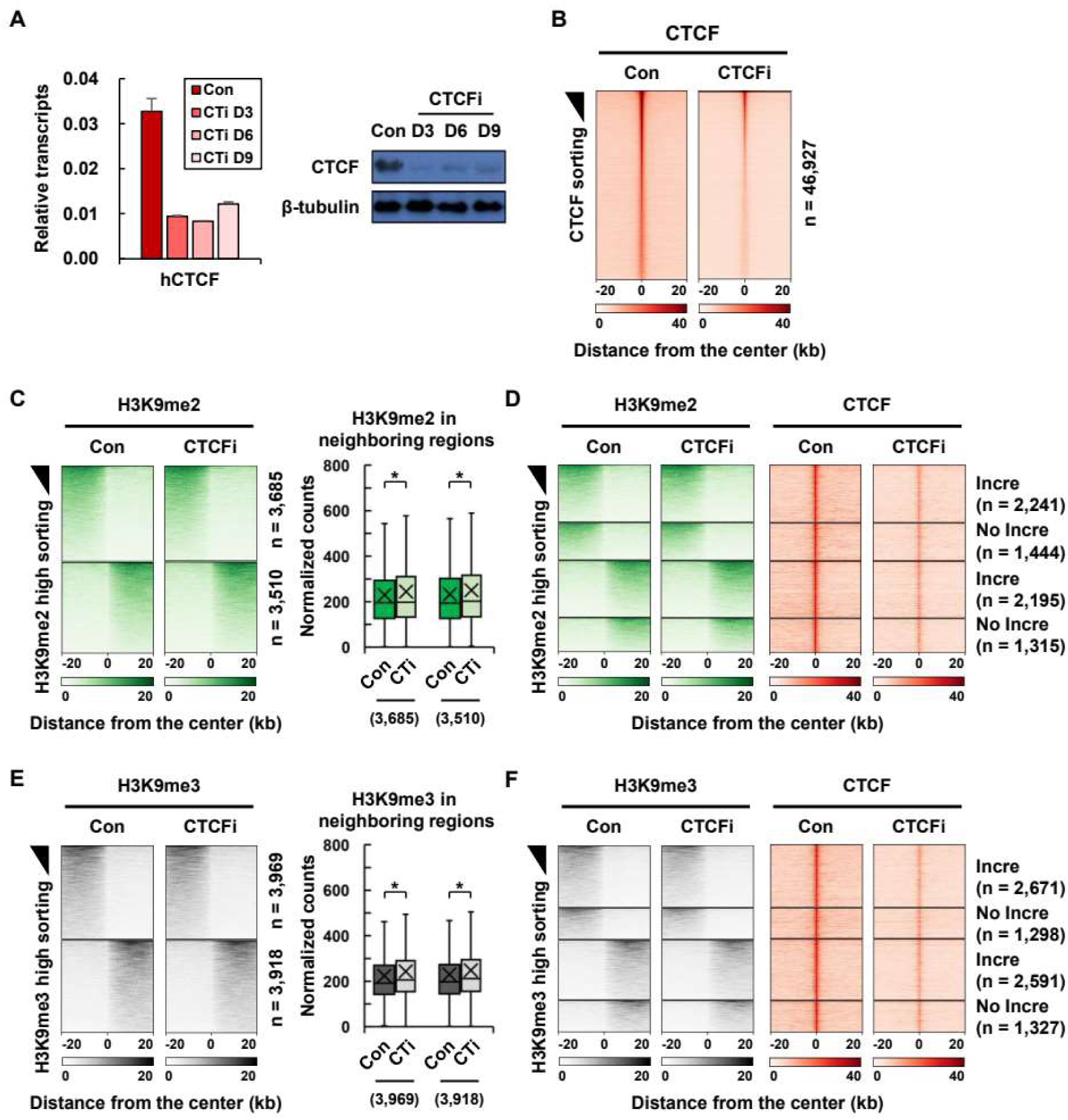
Spreading of H3K9me2 and H3K9me3 in CTCF-depleted K562 cells. (A) Expression levels of the CTCF gene were measured at the mRNA and protein levels using RT-qPCR and Western blot, respectively, in control (Con) and CTCF knockdown (CTCFi) K562 cells. (B) ChIP-seq signals of CTCF were visualized as heatmaps around its target binding sites in Con and CTCFi cells. (C) H3K9me2-enriched domains demarcated by CTCF-binding sites (n = 7,195) were identified based on a signal difference exceeding 100 and more than a 2-fold change between 20 kb upstream and downstream regions of each site. Heatmaps display H3K9me2 signals within these domains and their neighboring regions in Con and CTCFi cells. Based on genomic orientation, 3,685 of the H3K9me2- enriched domains are located upstream of CTCF-binding sites, and 3610 are located downstream. Normalized H3K9me2 signal counts in neighboring regions were quantified and presented as box plots. (D) H3K9me2-enriched domains were further classified based on whether H3K9me2 signals in neighboring regions increased (Incre) or did not increase (No Incre) in CTCFi cells. H3K9me2 and CTCF signals were visualized as heatmaps in Con and CTCFi cells for each group. (E, F) H3K9me3- enriched domains (n = 7,887) were defined based on a signal difference exceeding 200 and more than a 2-fold change between 20 kb upstream and downstream regions of each CTCF-binding site, and visualized using the same criteria as panels C and D. P-values were calculated using the Wilcoxon signed-rank test. *P<0.01

### Non-spreading boundaries of H3K9me2 and H3K9me3 are flanked by neighboring regions with a more active chromatin state

To investigate the relationship between the spreading of H3K9me2 and H3K9me3 and the underlying chromatin structure around CTCF-bound boundaries, we analyzed histone modifications in control K562 cells with intact CTCF binding. Figure 3A shows the enrichment patterns of H3K9me2, H3K9me3, and H3K9ac around boundaries with and without H3K9me2 spreading. For this analysis, H3K9me2-enriched domains were aligned upstream in a directional manner. The levels of H3K9me2 within these domains and their neighboring regions were comparable regardless of spreading, as shown by the profile plots (Fig. 3A). However, the neighboring regions adjacent to H3K9me2-enriched domains exhibited weaker H3K9me3 signals and stronger H3K9ac signals at boundaries where spreading did not occur, compared to those where spreading did occur. For H3K9me3-enriched domains, H3K9me3 levels within these domains and their neighboring regions were similar between spreading and non-spreading boundaries (Fig. 3B). Notably, neighboring regions flanking H3K9me3-enriched domains showed lower levels of H3K9me2 and higher levels of H3K9ac at non-spreading boundaries, relative to spreading boundaries. Together, these findings indicate that the non-spreading boundaries of H3K9me2 or H3K9me3-enriched domains are flanked by neighboring regions with a more active chromatin state than the spreading boundaries.

**Figure 3.**
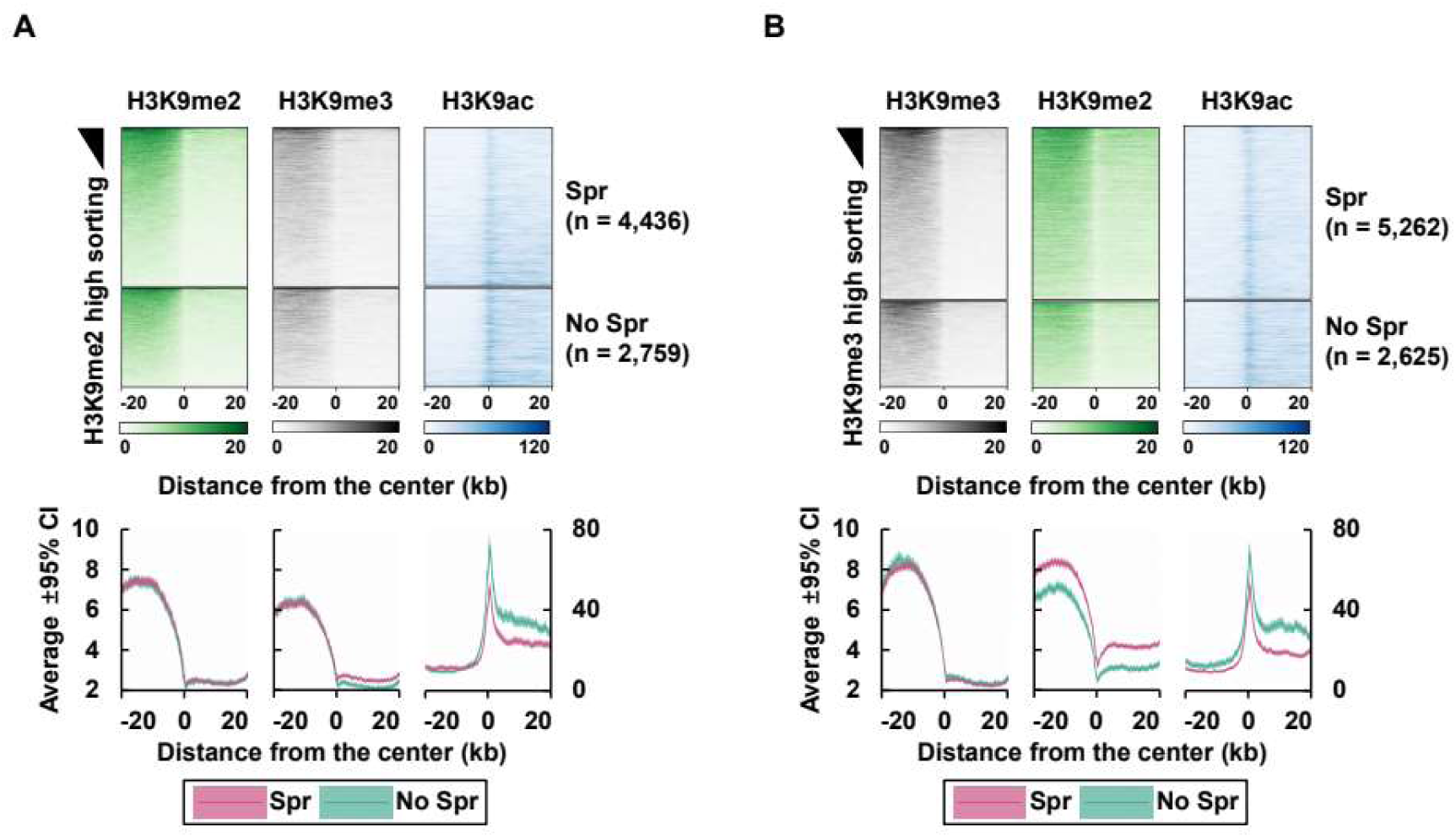
Chromatin structure at the spreading and non-spreading boundaries of H3K9me2 or H3K9me3. (A) ChIP-seq signals of H3K9me2, H3K9me3, and H3K9ac were visualized as heatmaps around H3K9me2-spreading boundaries (Spr, n = 4,436) and non-spreading boundaries (No Spr, n = 2,759) in control cells. (B) ChIP-seq signals of the same three histone modifications were visualized around H3K9me3-spreading boundaries (Spr, n = 5,262) and non-spreading boundaries (No Spr, n = 2,625). Profile plots below the heatmaps display average signal intensities with ±95% confidence intervals (CIs).

### Gene transcription is significantly reduced in neighboring regions where H3K9me2 or H3K9me3 spread

To examine the effect of H3K9me2 or H3K9me3 spreading on gene transcription, we performed RNA-seq in control and CTCFi K562 cells and measured transcription levels of annotated genes as CPM values using edgeR. We identified the TSSs of 1,292 genes with transcription levels over 1 CPM within 20 kb of neighboring regions where H3K9me2 had spread (Fig. 4A, B). When the transcription levels of these genes were compared, a significant reduction was observed in CTCFi cells compared to control cells (Fig. 4B). A similar reduction in transcription was observed in neighboring regions where H3K9me3 had spread (1,308 genes) (Fig. 4C). However, significant transcriptional changes were not detected in neighboring regions where the spreading of H3K9me2 or H3K9me3 did not occur (Supplemental Fig. S2). These results indicate that the spreading of either H3K9me2 or H3K9me3 leads to transcriptional repression. Representative genomic regions containing domains enriched with H3K9me2 or H3K9me3 are shown in Figure 4D and E, where the spreading of these modifications into adjacent regions upon CTCF loss is associated with transcriptional repression.

**Figure 4.**
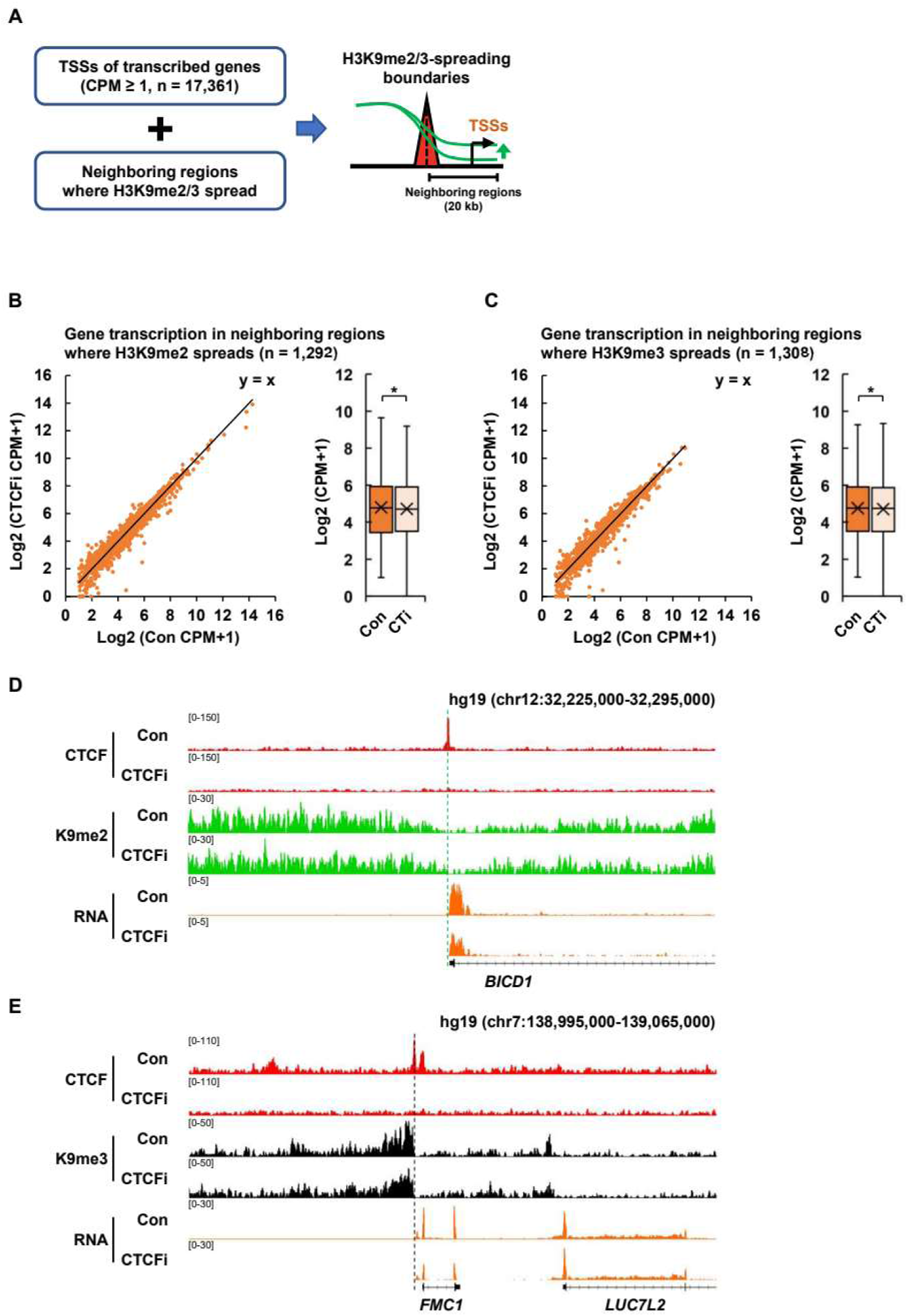
Gene transcription in neighboring regions where H3K9me2 or H3K9me3 spreads. (A) A schematic diagram illustrates the identification of genes located within neighboring regions where H3K9me2 or H3K9me3 spreads, based on the overlap between transcription start sites (TSSs) of transcribed genes (CPM ≥ 1) and these regions. Transcription levels of genes within regions where H3K9me2 (B) or H3K9me3 (C) spreads were visualized using scatter plots and box plots in control (Con) and CTCF knockdown (CTCFi) cells. Black diagonal lines in the scatter plots indicate a slope of 1. Genome browser views display ChIP-seq tracks for CTCF, H3K9me2, and H3K9me3, along with RNA-seq tracks, in Con and CTCFi cells across representative genomic regions containing H3K9me2- (D) or H3K9me3-enriched domains (E). Green and grey dash lines on the browsers represent the boundaries of domains enriched with H3K9me2 and H3K9me3, respectively. P-values were calculated using the Wilcoxon signed-rank test. *P<0.01

### Transcriptional reduction is specifically associated with the spread of H3K9me2 rather than H3K9me3 in comprehensive analysis

Since both H3K9me2 and H3K9me3 are markers of repressive chromatin and are often observed in the same genomic regions, it is unclear which modification specifically or primarily contributes to transcriptional repression in neighboring regions where these modifications have spread. Here, we overlapped the boundaries of H3K9me2-enriched domains with those of H3K9me3-enriched domains and identified 3,629 boundaries demarcating domains enriched with both modifications, accounting for about half of the boundaries of either H3K9m2- or H3K9me3-enriched domains (Fig. 5A). These boundaries were categorized into four groups based on the spreading patterns of H3K9me2 and H3K9me3 following CTCF depletion: both H3K9me2 and H3K9me3-spreading, H3K9me2-only spreading, H3K9me3-only spreading, and neither spreading. Although the most frequent cases were those in which both H3K9me2 and H3K9me3 spread together, these two modifications did not always spread concomitantly.

**Figure 5.**
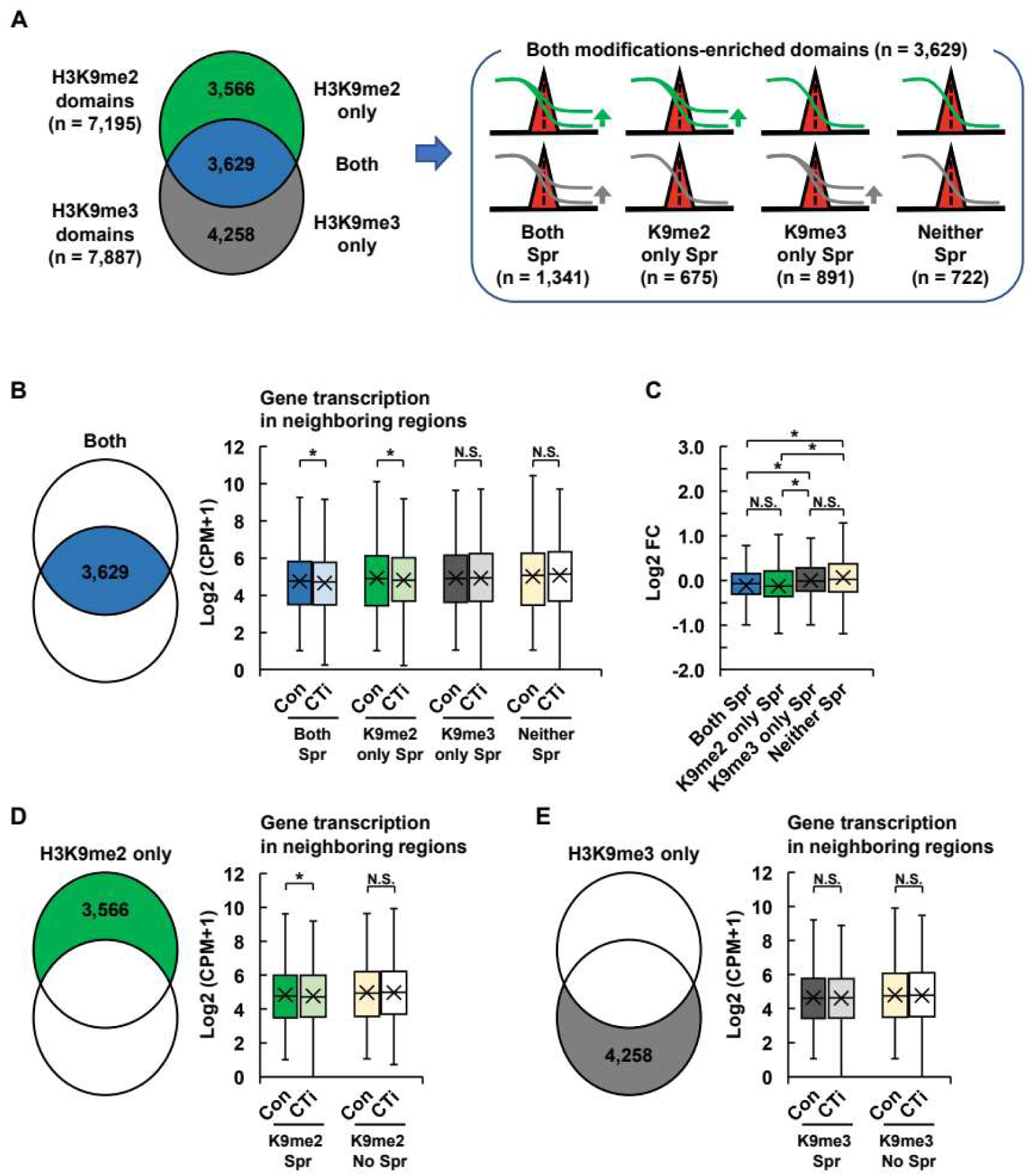
Gene transcription in neighboring regions of domains enriched with both H3K9me2 and H3K9me3 or either modification alone. (A) Boundaries of H3K9me2-enriched domains were overlapped with those of H3K9me3-enriched domains. Co-enriched domains with both modifications (n= 3,629) were classified into four groups based on the spreading pattern of H3K9me2 and H3K9me3 in CTCF knockdown (CTCFi) cells: both H3K9me2 and H3K9me3 spreading (Both Spr), H3K9me2- only spreading (K9me2 only Spr), H3K9me3-only spreading (K9me3 only Spr), and no spreading (Neither Spr). (B) Transcription levels of genes located in neighboring regions of each group (Both Spr: n = 455; K9me2 only Spr: n = 302; K9me3 only Spr: n = 368; Neither Spr: n = 300) were visualized using box plots in control (Con) and CTCFi cells. (C) Box plots show the log2 fold changes in transcription levels between Con and CTCFi cells for each group. (D, E) Domains enriched with either H3K9me2 or H3K9me3 alone were further divided based on whether spreading occurred in CTCFi cells. Box plots display transcription levels of genes located in neighboring regions of H3K9me2-only domains (K9me2 Spr: n = 652; K9me2 No Spr: n = 427) (D), and H3K9me3-only domains (K9me3 Spr: n = 597; K9me3 No Spr: n = 378) (E), in Con and CTCFi cells. P-values were calculated using the Wilcoxon signed-rank test (B, D, and E) and the Kruskal–Wallis test (C). NS: not significant; *P < 0.01

When gene transcription was compared between control and CTCFi cells, a significant reduction was observed in neighboring regions where either both H3K9me2 and H3K9me3 or only H3K9me2 had spread, but not in regions where only H3K9me3 or neither modification had spread (Fig. 5B). The degree of transcriptional reduction was not significantly different between regions where both modifications had spread and those where only H3K9me2 had spread, whereas a significant difference was observed between H3K9me2-only and H3K9me3-only spreading regions (Fig. 5C). This H3K9me2-dependent, but H3K9me3-independent, transcriptional repression was also observed in neighboring regions of domains enriched solely for H3K9me2 or H3K9me3 (Fig. 5D, E). Overall, this comprehensive analysis indicates that the spreading of H3K9me2, rather than H3K9me3, is primarily responsible for transcriptional repression.

## Discussion

Our study highlights the role of CTCF-binding sites as boundaries and barriers for chromatin domains enriched in H3K9me2 or H3K9me3 in mammalian cells (Fig. 6). Genome-wide analysis revealed that a subset of CTCF-binding sites demarcates these repressive domains and prevents the spreading of H3K9me2 and H3K9me3 into adjacent chromatin regions. In the absence of CTCF binding, such spreading appears to be constrained by neighboring regions of active chromatin state. Although reduced gene transcription was observed in regions where H3K9me2 or H3K9me3 had spread, comprehensive and comparative analysis indicates that this transcriptional repression is more strongly associated with the spreading of H3K9me2 than with that of H3K9me3.

**Figure 6.**
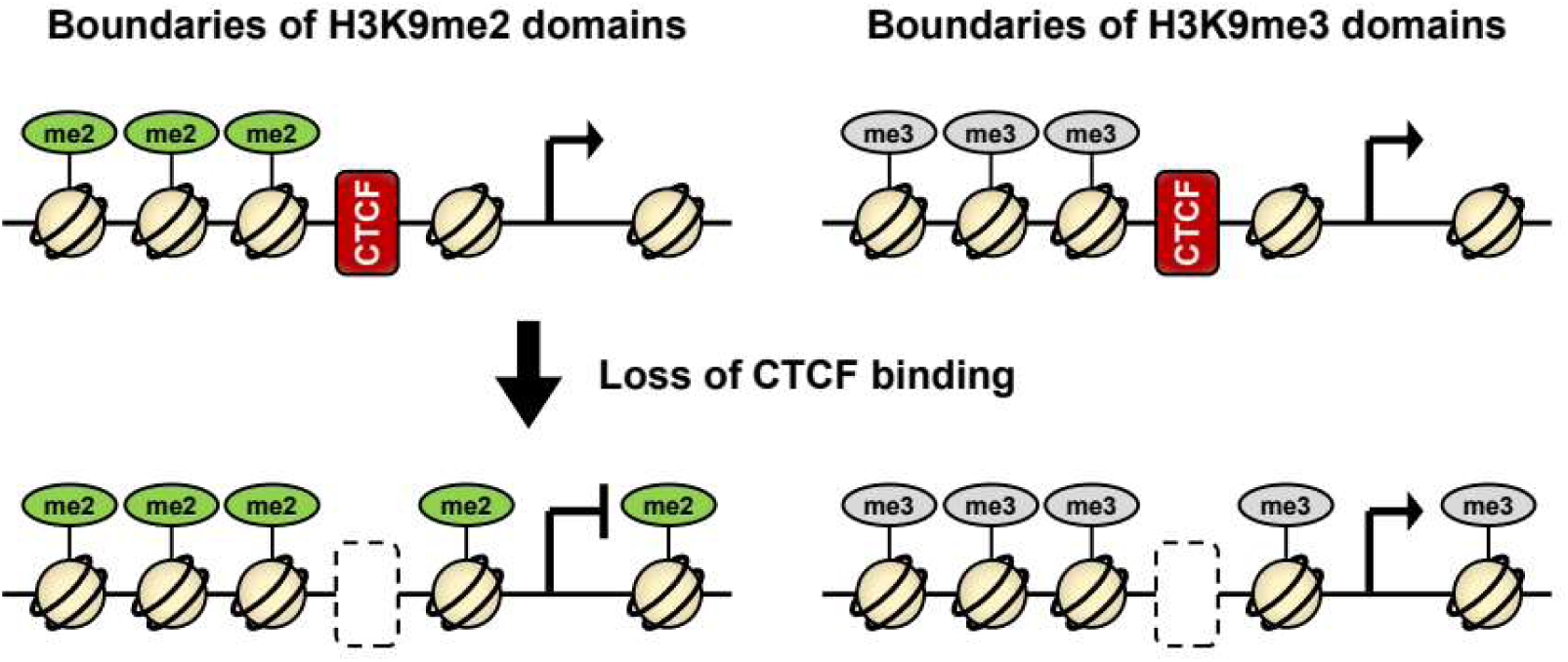
Role of CTCF-binding sites in regulating the spread of H3K9me2- and H3K9me3- enriched chromatin domains in human cells. CTCF-binding sites act as boundaries that demarcate chromatin domains enriched with H3K9me2 or H3K9me3 and function as barriers to prevent the spreading of these repressive histone modifications into neighboring regions in human K562 cells. Upon loss of CTCF binding, both H3K9me2 and H3K9me3 can extend beyond their original domains; however, only the spreading of H3K9me2 is associated with transcriptional repression of genes in the adjacent regions.

CTCF-binding sites have been reported to delimit chromatin domains enriched in several kinds of histone modifications, such as H3K27me2, H3K27me3, H3K4me3, or H3K27ac (Cuddapah et al. 2009; Chang et al. 2010; Kim et al. 2011a; Narendra et al. 2015; Kang et al. 2021). Their barrier activity has been suggested based on the spreading of these modifications following the loss of CTCF binding (Kim et al. 2011a; Narendra et al. 2015; Kang et al. 2021). Our genome-wide study revealed that CTCF-binding sites serve similar roles as boundaries and barriers in chromatin domains enriched with H3K9me2 or H3K9me3. The role of CTCF in restricting H3K9me3 spreading has been reported in *Drosophila*, where the mutations in two CTCF-binding motifs within the *Fab-8* boundary element led to the loss of its barrier activity at a homozygous transgenic locus, resulting in the spreading of repressive chromatin marked by H3K9me3 and H3K27me3 (Lu et al. 2018). In fission yeast, the LUC7 protein appears to play a similar role to CTCF by binding to heterochromatin borders (Wang et al. 2013). Expansion of H3K9me2 upon CTCF depletion has been observed at several gene loci in human epithelial cells (Jose et al. 2014). Taken together, CTCF-binding sites are thought to contribute to the demarcation of chromatin domains enriched with H3K9me2 or H3K9me3 and to prevent the spreading of these modifications from the domains into neighboring regions in mammalian cells.

The chromatin structure of neighboring regions appears to influence the spreading of H3K9me2 and H3K9me3 upon the loss of CTCF binding, as boundaries without the spreading were flanked by chromatin regions with higher levels of H3K9ac and lower levels of H3K9me2/3 in our analysis. The role of active chromatin in inhibiting the propagation of H3K9me2-enriched domains has been showed in fission yeast lacking LEO1, a component of PAF complex, where invasion of heterochromatin marked by H3K9me2 into adjacent euchromatin domains was accompanied by a reduction in Leo1- dependnent histone H4K16ac at boundary regions (Verrier et al. 2015). Furthermore, tethering a histone acetyltransferase to boundary chromatin suppressed the spreading of H3K9me2 in the absence of LEO1. A more direct role of H4K16ac was elucidated in the inverted repeats flanking heterochromatin at centromeres in fission yeast, where H4K16ac was recognized by the BDF2 protein, which is required for the boundary function of the inverted repeats (Wang et al. 2013). Loss of H4K16ac, either through histone acetyltransferase disruption or H4K16R mutation, promoted the spreading of H3K9me2. Similarly, when ectopic H3K9me3 was introduced at hundreds of genomic loci using a CRISPR-based system in human osteosarcoma cell line U2OS cells, this modification tended to spread into inactive chromatin but was largely restricted from active chromatin (Feng et al. 2020). Together, these findings suggest that an active chromatin state in neighboring regions can antagonize the spreading of H3K9me2 and H3K9me3 in a CTCF-independent manner.

In our study, we distinguished the spreading of H3K9me2 from that of H3K9me3 by separately analyzing chromatin domains enriched with both modifications, as well as those enriched with either modification alone. This analysis revealed that approximately half of the domains enriched with H3K9me2 or H3K9me3 were co-enriched with both marks, where these modifications often spread concomitantly. However, the spreading of each modification alone was also observed in some domains. These findings indicate that transcriptional repression of genes is primarily associated with the spreading of H3K9me2 rather than H3K9me3. This is consistent with a report showing that ectopically induced H3K9me3 is insufficient for gene silencing in a chromosome-wide study (Feng et al. 2020). Nevertheless, several studies have documented gene silencing accompanied by H3K9me3 at specific loci, although whether H3K9me2 was also involved remains unclear in these reports (Allan et al. 2012; Scarola et al. 2015). In our study, the spreading of H3K9me3 may have been too weak to repress gene transcription, or insufficient time may have passed to observe its effects. Taken together, our study provides a genome-wide, comparative analysis of H3K9me2 and H3K9me3 spreading following the loss of CTCF binding in mammalian cells and their respective impacts on gene transcription. Although the spreading occurred in both modifications after CTCF depletion, the propagation of H3K9me2 appears to play a more specific role in transcriptional repression.

## Methods

### Cell culture

K562 cells and 293FT cells were cultured in RPMI 1640 and DMEM media, respectively, each supplemented with 10% fetal bovine serum (FBS, Gibco). The DMEM medium for 293FT cells was additionally supplemented with non-essential amino acids (NEAA, Gibco). All cells were maintained at 37°C in a humidified incubator with 5% CO2.

### CTCF knockdown using lentiviral shRNA

The expression of *CTCF* gene in K562 cells was inhibited using TRC lentiviral short hairpin RNA (shRNA) vectors (Sigma-Aldrich, TRCN0000218498), as previously described (Kim et al. 2020). Lentiviral vectors containing CTCF shRNA gene were transfected into 293FT cells with the Virapower packaging mix (Invitrogen) using Lipofectamine 2000 (Invitrogen). After 72 hours (h), the viral supernatant was harvested and used to transduce K562 cells in the presence of 6 μg/mL polybrene. At 24 h after transduction, the cells were selected with 2 μg/mL puromycin. Empty shRNA vectors were used as an experimental control. *CTCF* expression was assessed at day 3, 6, and 9 after the transduction at the mRNA and protein levels using RT-qPCR and Western blot, respectively.

### Reverse transcription-qPCR (RT-qPCR)

Total RNA was extracted from K562 cells (2 × 10^6^) using QIAzol (Qiagen) and reverse-transcribed into cDNA using a GoScript kit (Promega) according to the manufacturer’s instructions. *CTCF* cDNA levels were quantified by qPCR and normalized to those of *ACTB*. Primer sequences used for this study are listed in Supplemental Table S1.

### Western blot

Proteins were extracted from K562 cells (2 × 10^6^) using RIPA buffer supplemented with protease inhibitors. Equal amounts of proteins were separated by electrophoresis on an 8% SDS-PAGE gel and transferred to a 0.2 μm nitrocellulose membrane. After blocking in PBST buffer containing 5% skim milk, the membrane was incubated overnight at 4°C with antibodies against CTCF (Millipore, 07-729) followed by incubation with an HRP-conjugated anti-rabbit secondary antibody (GeneTex, GTX213110-01) for 1 h at room temperature in PBST containing 1% skim milk. The antibody against β-tubulin was served as an experimental control. Protein signals were detected using an ECL reagent (Advansta).

### Chromatin immunoprecipitation (ChIP)

ChIP assays were performed as previously described (Kim et al. 2011b). Briefly, K562 cells (1×10^7^) were cross-linked with 1% formaldehyde. Chromatin was fragmented using 100 U of MNase and incubated with specific antibodies. Immunocomplexes were recovered using protein A agarose beads (Millipore). Immunoprecipitated DNA was purified by phenol extraction and ethanol precipitation. The antibodies used were CTCF (Millipore, 07-729), H3K9ac (Abcam, ab4441), H3K9me2 (Abcam, ab1220), and H3K9me3 (Abcam, ab8898). Spike-in chromatin (Active Motif; 25 ng for CTCF and 50 ng for the others) and spike-in antibody (Active Motif; 1 μg for CTCF and 2 μg for the others) were included as internal controls. ChIP assays for H3K9ac, H3K9me2, H3K9me3, and input were repeated with the spike-in controls (Supplemental Fig. S1).

### ChIP-seq library preparation

ChIPed DNA (0.5 ng for CTCF, 1 ng for H3K9ac, and 10 ng for H3K9me2 and H3K9me3) and input DNA (10 ng) were processed using the NEBNext Ultra II DNA Library Prep Kit (NEB) according to the manufacturer’s instructions. The DNA was end-repaired, ligated with NEBNext adaptors, size-selected at 175 bp using NEBNext Sample Purification Beads, and amplified with NEBNext Multiplex Oligos for Illumina. The libraries were quantified using a Qubit dsDNA HS Aassay Kit (Invitrogen) and sequenced as 100-base paired-end reads on an Illumina NovaSeq 6000 system.

### ChIP-seq data analysis

ChIP-seq raw reads were trimmed and filtered to remove low-quality reads using Trimmomatic (Bolger et al. 2014). Qualified reads were aligned to the hg19 canonical genome using Bowtie2 (Langmead and Salzberg 2012). Aligned BAM files were filtered by a minimum MAPQ quality score of 20 and sorted by chromosomal coordinates. Putative PCR duplicates were removed using RmDup (Li et al. 2009). CTCF-enriched peaks were identified with MACS2 using input data as controls (Feng et al. 2012). The amount of CTCF, H3K9ac, H3K9me2, and H3K9me3 was quantified by counting reads in the regions of interest using bedtools MultiCovBed (Quinlan and Hall 2010), followed by normalization with scaling factors to adjust for read counts aligned to the dm3 genome (spike-in control) and the hg19 genome (sample). To visualize ChIP-seq data, BAM files were converted to BigWig files using bamCoverage (Ramirez et al. 2016), with normalization based on the scaling factors. Heatmaps and profile plots were generated using computeMatrix, plotHeatmap, and ggplot2 (Wickham 2009; Ramirez et al. 2016). ChIP-seq signals along representative loci were visualized using the Integrated Genome Viewer (IGV) (Robinson et al. 2011).

### Boundary identification of chromatin domains enriched with H3K9me2 or H3K9me3

To identify CTCF-mediated boundaries of chromatin domains enriched with H3K9me2 or H3K9me3, signal intensities for these histone modifications were quantified in the 20 kb regions upstream and downstream of CTCF-enriched summits (n = 46,927). Normalized read counts were obtained by multiplying each raw read count by the corresponding scaling factor. Differences and fold-changes in H3K9me2 and H3K9me3 were calculated by comparing the normalized read counts between the upstream and downstream regions of CTCF-binding sites. Boundaries were defined as sites exhibiting a minimum difference of 100 and a fold change of at least 2 for H3K9me2, and a minimum difference of 200 and a fold change of at least 2 for H3K9me3, indicating a sharp transition in histone modification enrichment.

### RNA-seq library preparation

Total RNA was extracted using the RNeasy Plus Mini Kit (Qiagen) and assessed for quality using the Qubit RNA IQ Kit (Invitrogen), with RNA IQ values greater than 7. Ribosomal RNAs (rRNAs) were removed using the RiboMinus Human/Mouse Module (Invitrogen), and 125 ng of rRNA-depleted RNA was used to construct libraries with the NEBNext Ultra II Directional RNA Library Prep Kit (NEB), following the manufacturer’s instructions. The procedure included RNA fragmentation, priming with random primers, first-strand cDNA synthesis incorporating dUTP, second-strand cDNA synthesis, end repair, adaptor ligation, PCR amplification for 8 cycles using adaptor-specific primers, and bead-based purification. Final libraries were quantified using the Qubit dsDNA HS Assay Kit (Invitrogen) and sequenced as 100-base paired-end reads on an Illumina NovaSeq 6000 system.

### RNA-seq data analysis

Raw RNA-seq reads were trimmed and filtered to remove low-quality sequences using Trimmomatic (Bolger et al. 2014). Filtered reads were aligned to the hg19 reference genome using STAR with an exon annotation GTF file from the UCSC database (Dobin and Gingeras 2015). Gene-level read counts were obtained using featureCounts for all annotated genes (Liao et al. 2014), and transcription levels were calculated as counts per million (CPM) using edgeR (Robinson et al. 2010). TSSs were identified using a gene BED file from the UCSC database, and a BED file containing TSSs plus 1 bp toward gene bodies was used to identify genes located in neighboring regions of H3K9me2 or H3K9me3-enriched domains. For visualization, BigWig files were generated from aligned BAM files using bamCoverage, normalized to the effective genome size of hg19 (Ramirez et al. 2016). The RNA-seq data for control K562 cells were available in NCBI Gene Expression Omnibus (GEO) under accession number GSE211316.

### Statistical analysis

P-values were calculated using the Mann-Whitney U test or Kruskal-Wallis test for comparisons of independent datasets with unequal sample sizes. For paired datasets with equal sample sizes, p-values were calculated using the Wilcoxon signed-rank test or the Kruskal-Wallis test. The normality of ChIP-seq and RNA-seq data distribution was assessed using the Kolmogorov-Smirnov test.

### Data access

All raw and processed sequencing data generated in this study have been submitted to the NCBI Gene Expression Omnibus (GEO; https://www.ncbi.nlm.nih.gov/geo/) under accession number GSE298130.

## Acknowledgements

This research was supported by Basic Science Research Program through the National Research Foundation of Korea (NRF) funded by the Ministry of Education (NRF-2020R1I1A3054808) and by the National Research Foundation of Korea (NRF) grant funded by the Korea government (MSIT) (No. 2022R1C1C2006355).

